# Conserved and divergent peptide substrate binding properties of bacterial Hsp70

**DOI:** 10.1101/2022.08.13.503765

**Authors:** Yiyue Sun, Hongke Xu, Jiong Li, Hanmo Zhu, Hongwei Ma, Yiming Ma, Jiao Yang

## Abstract

The 70-kDa heat shock protein (Hsp70) interacts with the polypeptide segments of abundant native proteins to fulfill various cellular activities in both stress and normal conditions. However, a non-native linear polypeptide NR (NRLLLTG) is widely used for the study of Hsp70 substrate binding properties, which is too simple to reflect the complex status of Hsp70 substrates in living organisms. To further broaden our knowledge in this area, we screened 2645 polypeptides derived from 78 biologically relevant proteins and identified eight native peptide substrates (named VP1∼VP8) for bacterial Hsp70 DnaK. Consistent with previous findings, the amino acid distribution in VP1∼VP8 were enriched in aliphatic and basic residues, and most of their residues were buried in folded proteins as well. Besides, the substrate binding properties for seven polypeptides were largely the same as observed in NR, suggesting their conserved binding mode to DnaK. However, VP5, which contains more percentage of positively charged residues, demonstrates divergent substrate binding properties during *in*-*vitro* biochemical studies. Moreover, VP5 efficiently inhibits the refolding activity of DnaK and bacterial viability, implying its potential to be a good lead peptide for antibacterial drug development.

## Introduction

Protein homeostasis (proteostasis) network safeguards cellular proteome through coordinated protein synthesis, assembly/disassembly, folding, posttranslational modification, trafficking, localization, quality control and degradation.^1-2^ Cells have evolved a series of strategies to maintain a healthy proteostasis. Molecular chaperones are one of these strategies; in particular, 70-kDa heat shock protein (Hsp70) serves as a master player in maintaining proteostasis.^2-5^ During stress, Hsp70s bind with the polypeptide segments of misfolded or aggregated proteins to protect them from further denaturation until the stress is relieved.^4, 6-7^ Consistent with these properties, Hsp70s have been implicated in a wide spectrum of diseases such as cancer, neurodegenerative diseases, inflammation, and infectious diseases.^8-12^

All Hsp70 proteins consist of two highly conserved domains: an N-terminal nucleotide-binding domain (NBD) which exhibits ATPase activity, and a C-terminal substrate-binding domain (SBD, consisting of SBDα and SBDβ) which mainly binds to client peptide substrates.^5, 13-14^ Hsp70 chaperone activity strictly depends on the allosteric coupling of the NBD and SBD.^15-17^ Upon binding of ATP to the NBD, the substrate-binding pocket in SBDβ is widely open, giving a low affinity and fast exchange rate for the peptide substrates.^14-15, 18^ The binding of peptide substrates in the substrate-binding pocket can, in turn, accelerates the hydrolysis of ATP. When the NBD is occupied by ADP, the substrate-binding pocket is covered by SBDα and locked into a closed conformation, thus resulting in slower substrate binding kinetics and much higher binding affinity compared with the ATP-bound state.^17, 19-20^ However, the intrinsic ATPase rate of Hsp70 is low^18^ and requires co-chaperones J-domain proteins (JDPs; target and trap substrates via ATP hydrolysis)^20-21^ and nucleotide exchange factors (NEFs; accelerate ADP–ATP exchange)^20, 22^ to facilitate the working cycle.^5, 17, 23^

Peptide substrate has been widely accepted as the important tool to investigate the allosteric coupling mechanism of Hsp70.^6, 23-26^ So far, the well characterized peptide substrate for Hsp70s is a non-native linear short polypeptide NR (NRLLLTG).^19, 27^ and most recent progresses in interpreting Hsp70 allosteric coupling are fulfilled by using NR polypeptide.^28-30^ Recently, TRP2 (VYDFFVWLHYY), a polypeptide derived from the sequence of a tumor antigen, has been identified as a novel peptide substrate for *E. coli* Hsp70 DnaK.^31^ Interestingly, unlike the canonical binding mode of most identified polypeptides, TRP2 binds to an unexpected second substrate binding site of DnaK and exhibits different binding properties. This unexpected finding implies that there may exist distinct peptide binding properties for native protein-originated polypeptides (native polypeptides) in the process of allosteric coupling due to the versatility of native protein *in vivo*, but unfortunately till now very limited effort has been made in this field.

To further explore the substrate binding properties for native polypeptides of Hsp70s, we carried out a screening to identify DnaK binding polypeptides through an iPDMS-based peptide library which is composed of polypeptides derived from native proteins in this study. The sequence characteristics of the identified polypeptides were analyzed and the substrate binding properties for each native polypeptide were further investigated.

## Results

### iPDMS-based peptide microarray and the scheme for experimental design

In the present study, we utilized polymer coated initiator integrated poly (dimethysiloxane) (iPDMS) nanomembrane for the establishment of a peptide microarray consisting of 2645 polypeptides derived from native proteins, and most of these proteins are related to Hsp70 functions (Fig. 1A and Table S1). iPDMS nanomembrane is a novel solid supporting material which is able to prevent nonspecific protein adsorption to an “absolute” zero level thus to achieve high detection sensitivity,^32^ increasing the chance to identify more potential target protein-binding polypeptides (Fig. 1B∼1D). The principle of peptide microarray screening is based on indirect ELISA (Fig. 1E). And the microarray image was finally displayed in chemiluminescent signals (Fig. 1F).

**Figure 1.**
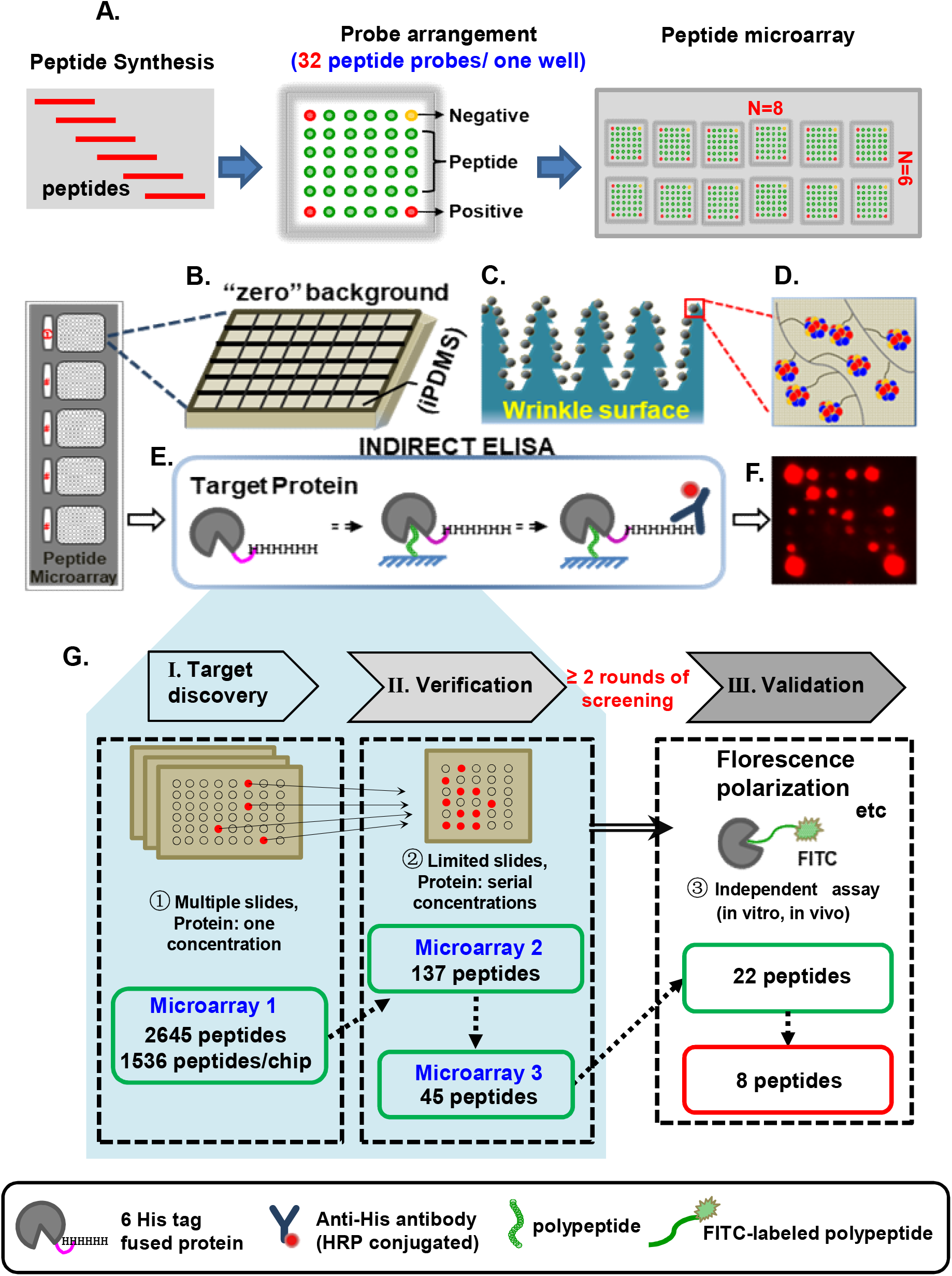
Schematic depiction of the iPDMS-based peptide microarray screening for discovery DnaK-binding polypeptides. **(A)** Polypeptides derived from native proteins were synthesized and immobilized onto chips. For each well, three positive spots, one negative spot and 32 polypeptide probes were immobilized. Finally, the peptide microarray was fabricated with 48 wells for each individual chip. **(B)** Peptide microarray was established by immobilizing polypeptides on the “absolute” zero background polymer coated initiator integrated poly(dimethysiloxane) (iPDMS) nanomembrane. **(C∼D)** iPDMS has a wrinkle surface which can increase the binding amount of the substance to be immobilized. **(E∼F)** To further identify polypeptides that can bind to target protein, peptide microarray will be incubated with target protein at first, then polypeptides with binding activities will bind to target protein. Protein/polypeptide complex will be further identified by HRP-conjugated antibody which specifically recognizes protein-fused tag and the chemiluminescent signals will be detected. **(G)** The screening process included three steps, (I) target discovery, (II) verification and (III) validation. In step III (Validation), polypeptides verified in Step II were further validated and investigated using independent assays (Florescence Polarization assay, etc.).

In order to efficiently identify and investigate the substrate binding properties for novel DnaK interacting polypeptides, three-step screening was introduced here (Fig. 1G): (i) Microarray 1 was screened against one DnaK concentration to initially acquire polypeptides which could interact with DnaK, so as to downsize peptide microarray for further screening (Step I. Target discovery); (ii) Both Microarray 2 and Microarray 3 were screened against multiple indicated concentrations of DnaK to exclude randomly responding polypeptides and increase the accuracy of our experiment (Step II. Verification); (iii) the investigations on substrate binding properties for identified interesting polypeptides through independent *in vitro* and *in vivo* assays (Step III. Validation).

### Polypeptides VP1∼VP8 showed DnaK concentration-dependent binding activities

According to the above screening strategy, 137 polypeptides were identified with DnaK-binding activities in the primary screening (Fig. 1G, Fig. 2A), showing a high response rate (∼5.2%). To exclude randomly responding hits here, the 137 hits were further incubated with DnaK protein at various concentrations (20, 10, 5, 2.5 and 1.25 μM), thus a large number of chemiluminescent signal values could be collected to plot a heat map (Fig. 2B). 45 polypeptides which showed excellent dose-dependent responses were performed for one more round screening with multiple concentrations of DnaK, in order to repeat their behaviors (Fig. 1G). Based on the Step II procedure, we obtained 22 Verified Polypeptides (VPs) with good DnaK concentration-dependent binding activities, of which, VP1∼VP8 showed lasting and strong chemiluminescent signals (Fig. 2C), implying their high binding affinities with DnaK. The detailed information for VP1∼VP8, including amino acid sequences and their corresponding native protein sources, were summarized in Fig. 2D.

**Figure 2.**
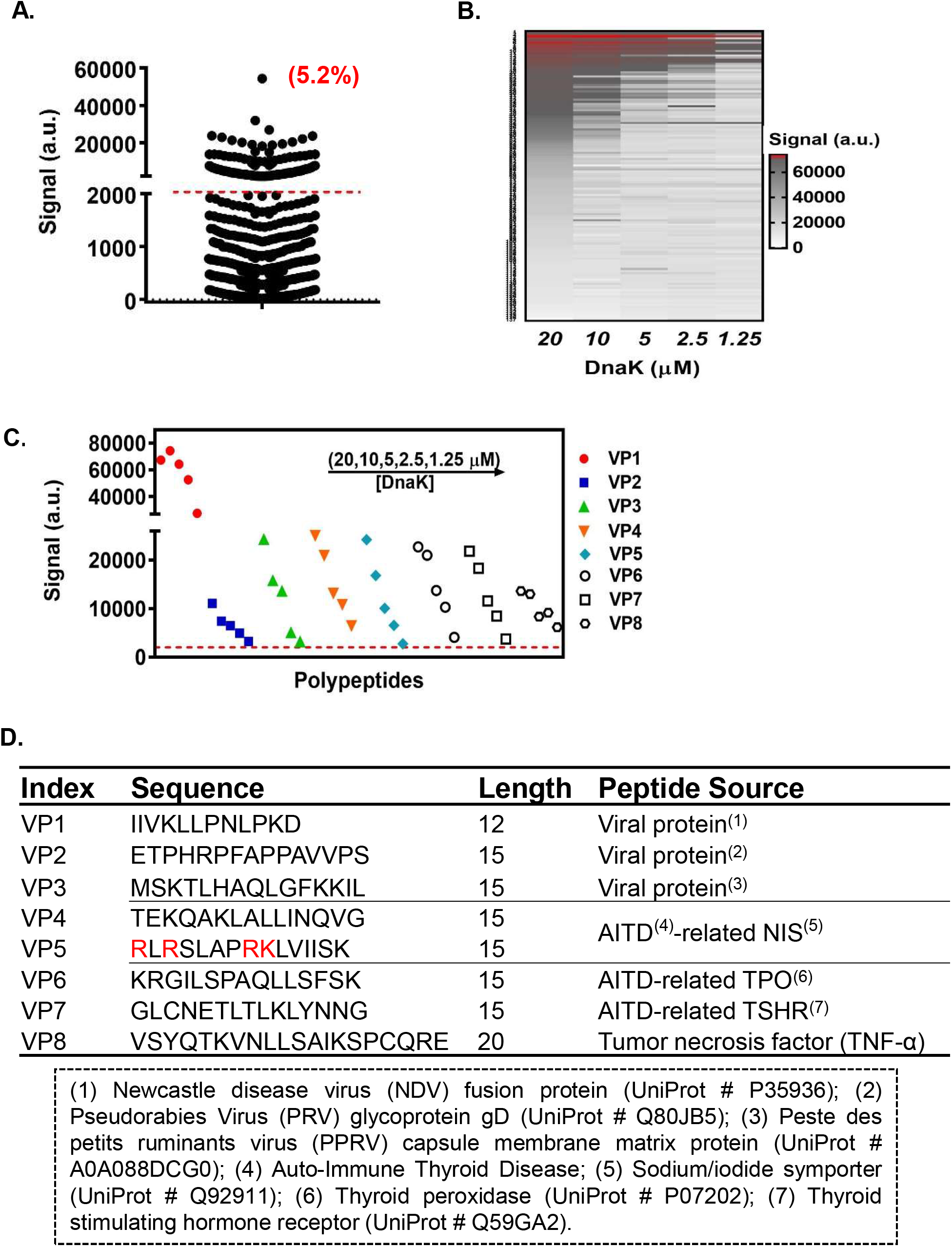
VP1∼VP8 were identified with DnaK binding activity. **(A)** In the high-efficiency screening, 5.2% polypeptides showed significant binding activities with 10 μM DnaK. Signal 2000 arbitrary units (a.u.) was set as the threshold (dashed line). **(B)** Heat map represented mean values of chemiluminescent signals in two independent experiments was used to analyze binding activities of polypeptides with various doses of DnaK (20, 10, 5, 2.5 and 1.25 μM, respectively). **(C)** VP1∼VP8 showed good chemiluminescent signals at five different concentrations of DnaK. **(D)** The amino acid sequences and the corresponding native proteins for VP1∼VP8.

### Sequence characterizations of VP1∼VP8 and their localizations within native protein structures

DnaK binding motif usually consists of a hydrophobic core of four to six residues enriched particularly in Leu, Ile, Val and Pro, which are mostly buried in folded proteins, flanked by positively charged residues Arg and Lys.^33-35^ At first, we tried to figure out if amino acid distribution in VP1∼VP8 showed the same characteristics as DnaK binding motif in native proteins. Consistent with previous findings, the residues Ile, Leu, Val, Pro, Arg and Lys were frequently shown in polypeptides VP1∼VP8, while residues Met, Cys and Asp were shown in a low occurrence (Fig. 3A).

**Figure 3.**
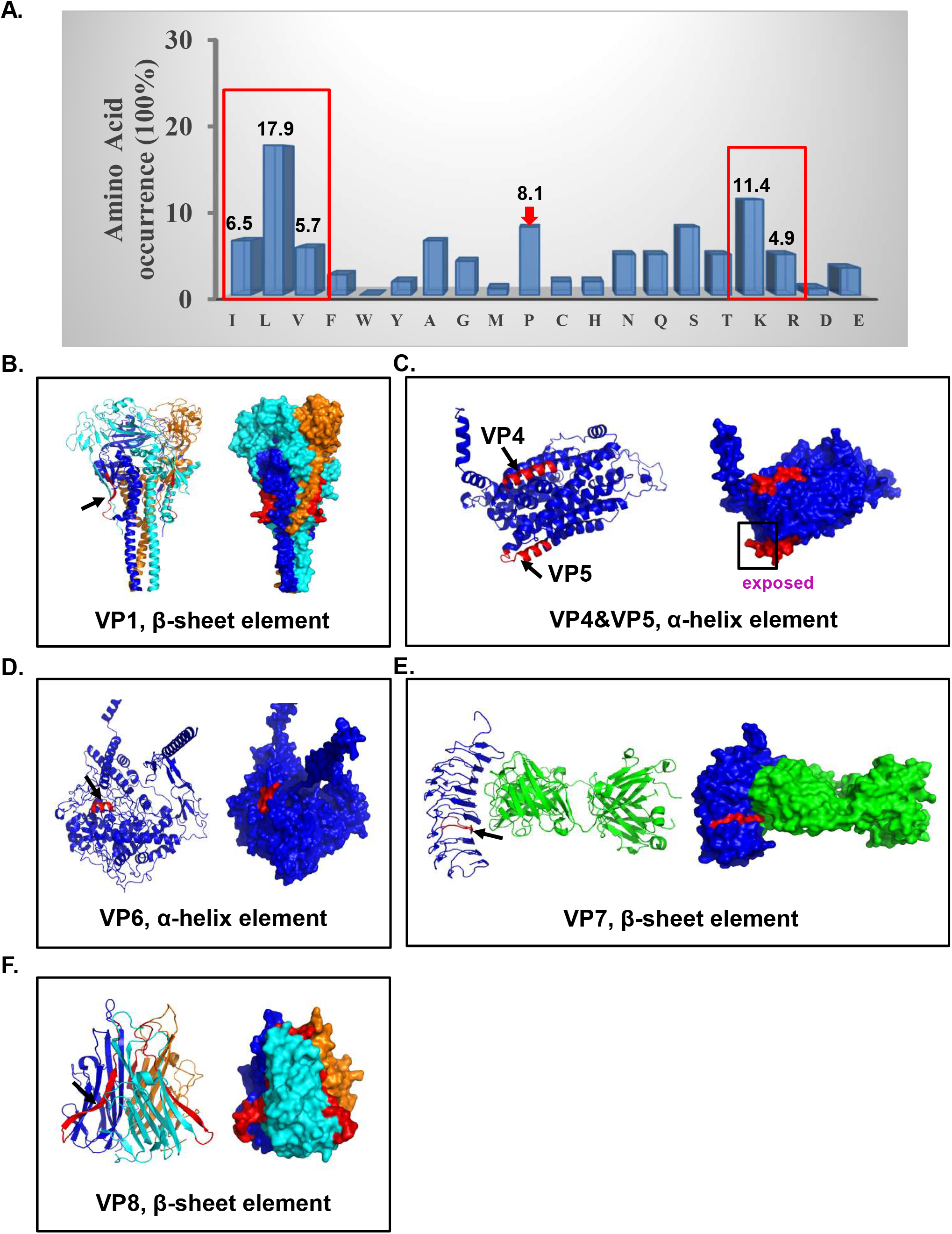
Analysis of the polypeptide sequences recognized by DnaK. **(A)** For polypeptides VP1∼VP8, the occurrence (%) of Ile, Leu, Val, Lys, Arg and Pro (indicated with red frame and arrow) was determined and shown on the top of each column, respectively. **(B-F)** Ribbon diagram and surface presentations of polypeptides corresponding native protein structures. The PDB IDs or AlphaFold model IDs were 1G5G (NDV fusion protein), AF-Q92911-F1 (human NIS), AF-P07202-F1 (human TPO), 2XWT (human TSHR) and 1TNF (human TNF-α), respectively. The location of VP1, VP4∼VP8 in corresponding native proteins was respectively highlighted in red.

Thanks to the availability of the corresponding three-dimensional protein structures, we were able to determine the localizations of polypeptides VP1 and VP4∼VP8 within their native protein structures, respectively. Highly consistent with previous reports,^33-35^ most residues of VP1 and VP8 are completely buried in trimer interface of NDV fusion protein (Fig. 3B) and human TNF-α (Fig. 3F). For other cases, though the side chains of residues in polypeptides are exposed at the protein surface, their peptide backbone are not extensively surface exposed (VP4 in Fig. 3C, Fig. 3D, Fig. 3E). More strikingly, we found that the N-terminal residues of VP5 (the verified DnaK binding sites in predicted structure), containing more percentage of positively charged amino acids than in regular DnaK binding motif (Fig. 2D), presented disordered conformation of exposed side chains and backbone (VP5 in Fig. 3C) within its native protein structure, divergent from most previously reported DnaK-binding polypeptides.

Moreover, our polypeptides are either located in (nearby) segments corresponding to β-sheet in native proteins (VP1 in Fig. 3B, VP7 in Fig. 3E, VP8 in Fig. 3F) or to α-helix in native proteins (VP4& VP5 in Fig. 3C, VP6 in Fig. 3D), similar to previous data as well.

### Conserved and divergent peptide substrate binding properties among VP1∼VP8

To illustrate how VP1∼VP8 binds to DnaK, the binding affinities of VP1∼VP8 to DnaK protein in the absence of ATP were evaluated first using fluorescence polarization assay (Fig. 4A and Fig. S1). VP1 and VP5 showed the best binding affinities among VP1∼VP8, with dissociation constants (K_*d*_) values a bit better than that of NR (Fig. 4A).

**Figure 4.**
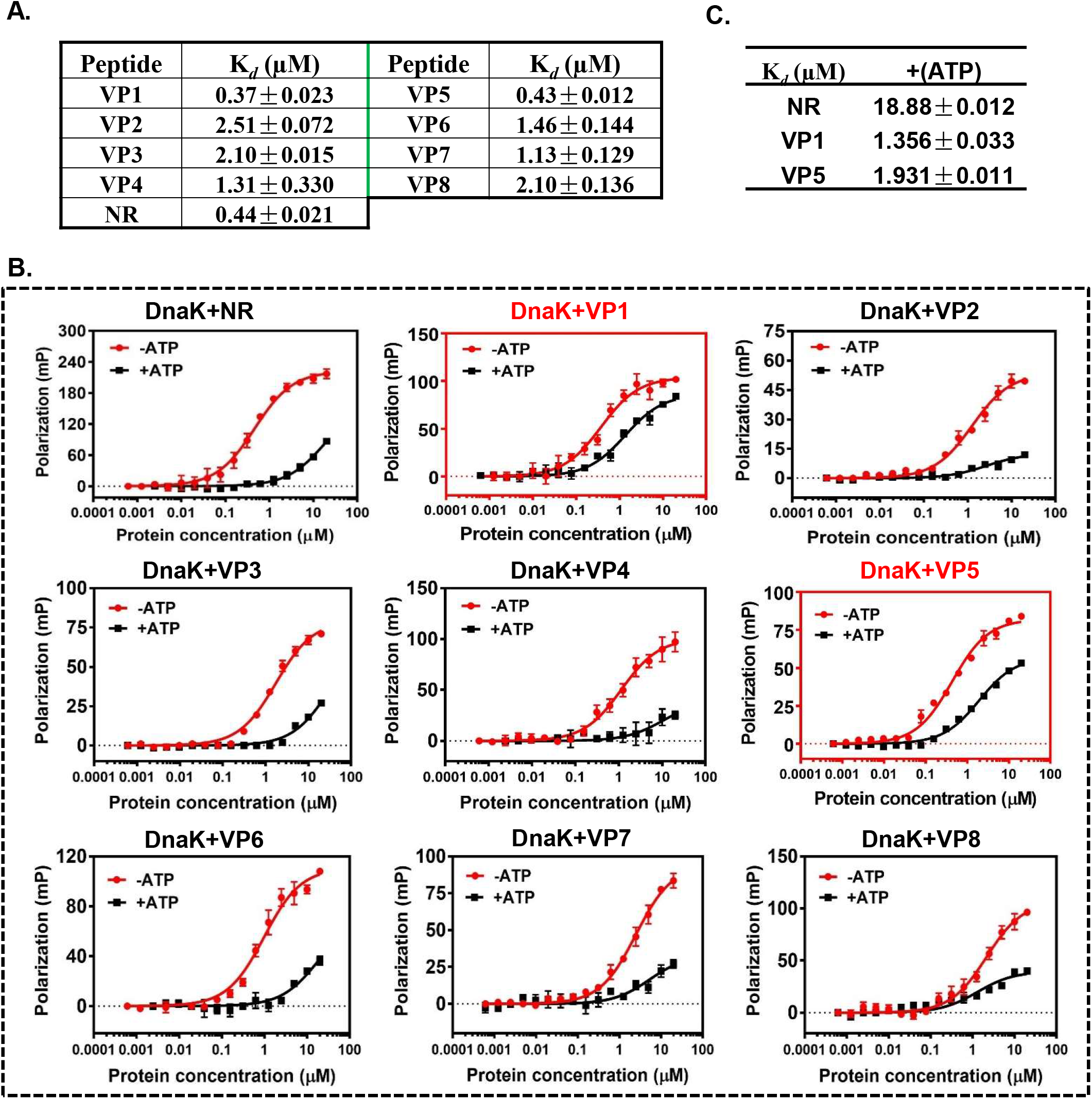
The substrate binding properties of VP1∼VP8. **(A)** The dissociation constants (K_*d*_) of VP1∼VP8 binding to DnaK were determined by fluorescence polarization assay. **(B)** The peptide substrate binding of VP1∼VP8 to DnaK in the absence of and presence of ATP. Similar to NR, VP2/3/4/6/7/8 showed conserved binding properties; while both VP1 and VP5 showed divergent binding properties, with limited signal decreases in fluorescence polarization assay at indicated protein concentrations in the presence of ATP. **(C)** The peptide binding affinity (K_*d*_) for NR, VP1 and VP5 to DnaK in the presence of ATP.

Consistent with previous studies, we observed that when DnaK bound with ATP, the binding affinity of DnaK for VP2/3/4/6/7/8 polypeptide was significantly reduced as observed in NR, suggesting their conserved and classical substrate binding properties (Fig. 4B). Interestingly, the affinity reduction of DnaK for VP1 or VP5 were not as obvious as that for NR in the presence of ATP (Fig. 4C), resulting in the observed divergent substrate binding properties (Fig. 4B).

We found the much slower binding rate of NR and most of the identified polypeptides (VP2/3/4/6/7/8) in the absence of ATP than that in the presence of ATP (Fig. 5A, Fig. S2), further implying the occurrence of classical allosteric coupling. Differently, whether in the presence of ATP or not, VP1 showed much slower binding kinetics than NR (Fig. 5B), thus its binding affinity to DnaK increased about 14-fold than NR when in the presence of ATP (Fig. 4C). Surprisingly, unlike NR and VP1, VP5 showed similar binding rate either in the absence of or presence of ATP (Fig. 5C), further suggesting its special binding mode to DnaK protein.

**Figure 5.**
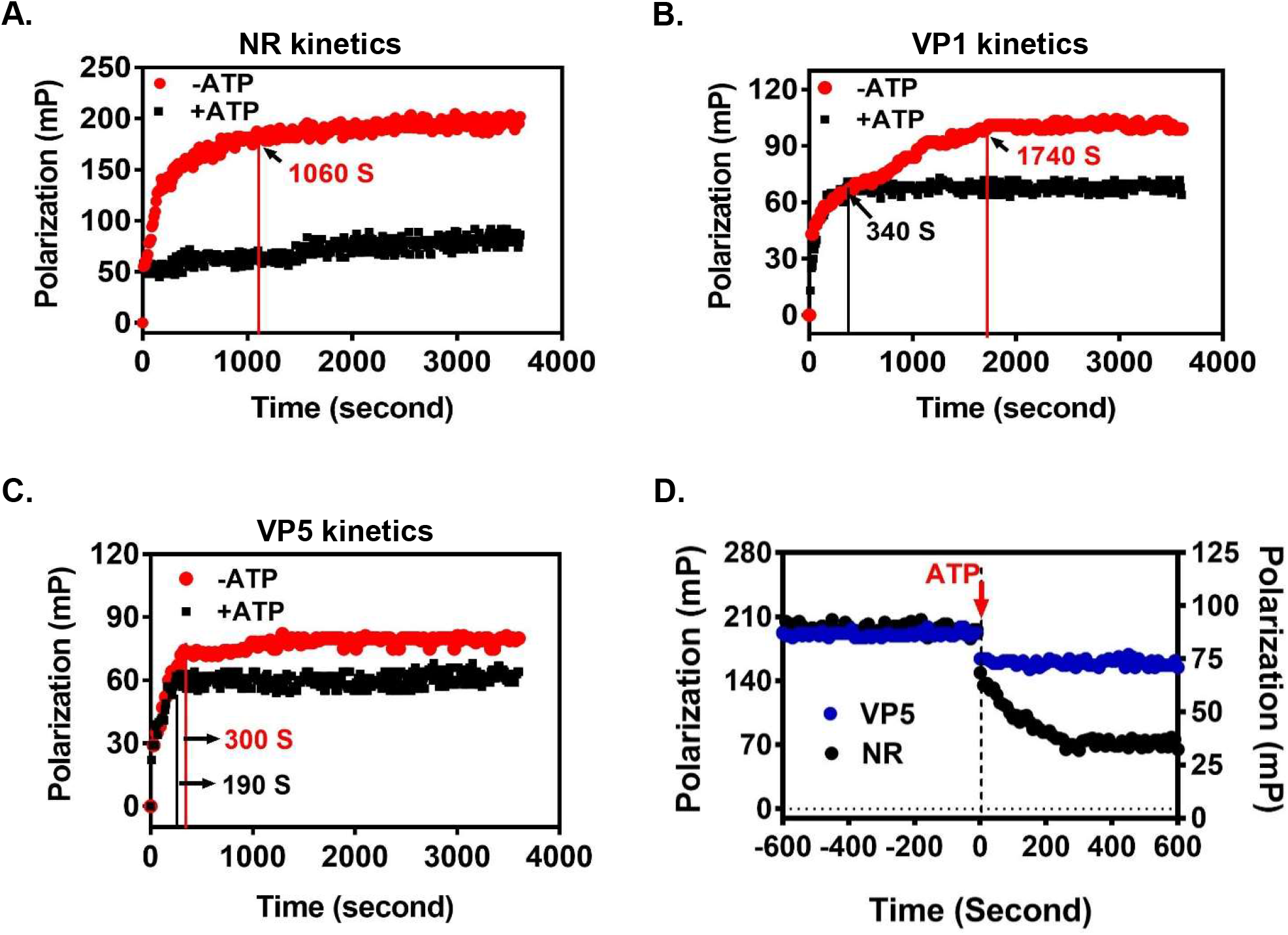
The unique property of substrate binding kinetics for VP5 in the absence of and presence of ATP. To dissect the divergent behaviors for polypeptides (VP1&VP5) in fluorescence polarization assay, further peptide binding kinetics were tested for NR **(A)**, VP1 **(B)** and VP5 **(C)** in the absence of and presence of ATP. The time for different peptide substrates binding to DnaK and getting equilibrium in the absence of ATP is ∼1060 s (NR), ∼1740 s (VP1) and ∼300 s (VP5), respectively. While, the equilibrium time for polypeptides (NR and VP1) binding to DnaK in the presence of ATP was much shorter and indicated with black arrow. **(D)** ATP obviously mediated NR release from DnaK (black dots, left y-axis), implying the occurrence of allosteric coupling; while the addition of ATP just mediated a little amount of VP5 release from DnaK (blue dots, right y-axis).

We further tested if adding ATP can cause the release of peptide substrates, and found that after adding ATP, NR or VP1/2/3/4/6/7/8 was able to release from DnaK, implying the regulation of substrate binding affinity by allosteric coupling mechanism (Fig. 5D, Fig. S3). However, adding ATP did not cause a significant release of VP5, indicating that VP5 binding affinity was not tightly regulated by classical allosteric coupling (Fig. 5D).

Till now, we concluded that VP1 showed divergent substrate binding property only in peptide substrate binding assay and its substrate binding affinity is still regulated by classical allosteric coupling. VP5 showed obviously divergent substrate binding properties during the whole process of our biochemical assays as mentioned above, and might bind to DnaK in a unique mode.

### VP5 disrupts the refolding activity of DnaK and inhibits bacterial growth more efficiently than NR

To evaluate if the binding of VP5 to DnaK has the potential to disturb the chaperone activity, we carried out the luciferase refolding assay to assess its performance. Compared with NR and most of polypeptides identified in this study, VP5 showed much stronger inhibitory effect on refolding activity (Fig. 6A, Fig. S4), but just less than VP4 polypeptide which may influence chaperone refolding activity through regulating co-chaperone activitiy of DnaJ or GrpE and needs further examinations in the future.

**Figure 6.**
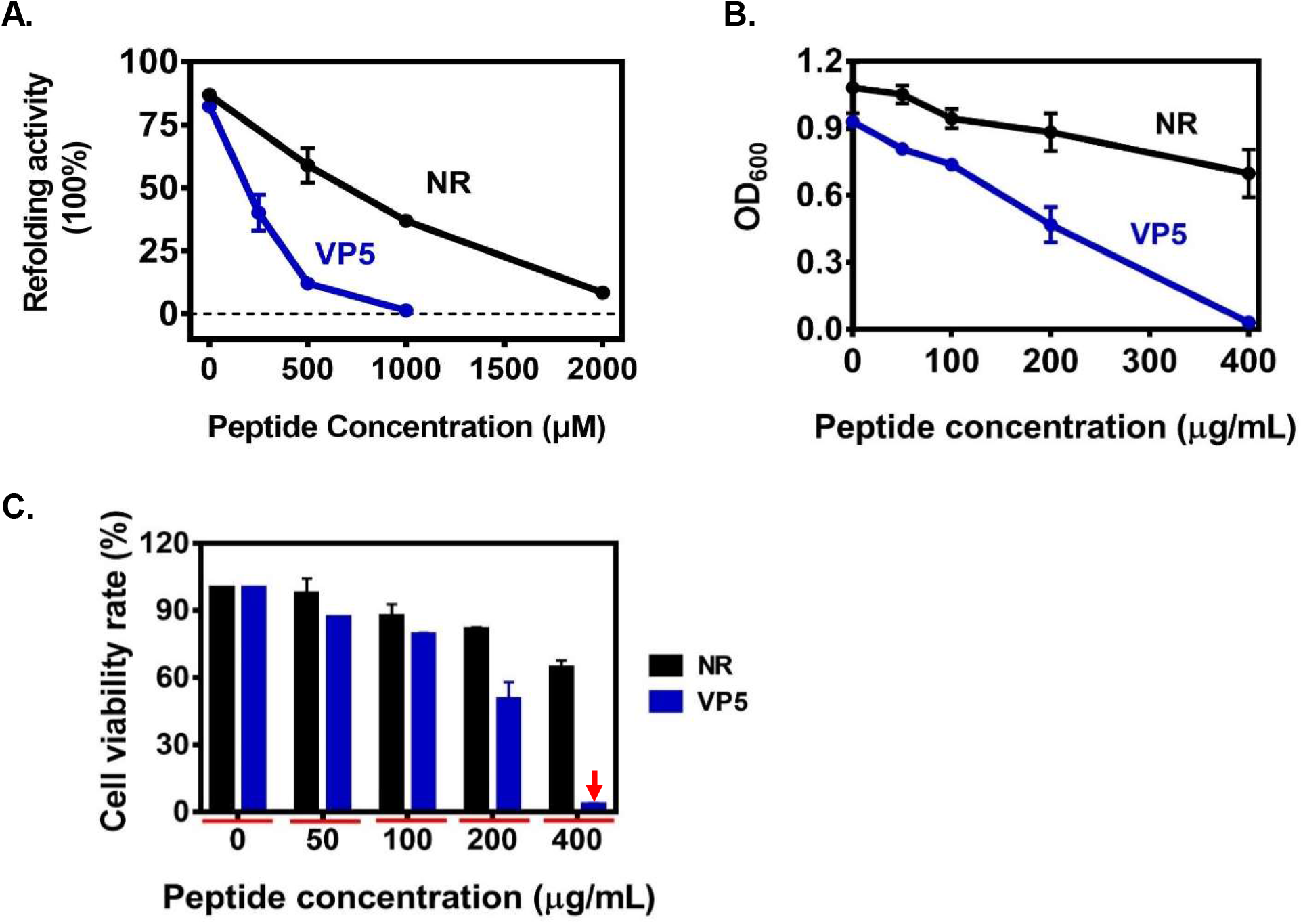
VP5 inhibits the DnaK-mediated refolding of luciferase and cellular viability. **(A)** VP5 inhibits the refolding activity of DnaK in a dose-dependent manner, and more efficiently than NR. **(B and C)** VP5 showed more efficient inhibitory activity against wild-type *E. coli* (ATCC 25922) cells than NR. Wild-type *E. coli* cells were seeded into 96-well plate. Serially diluted stock solutions of the polypeptides were then added into each well prior to overnight growth for ∼18 h at 42°C. **(B)** The OD_600_ of each well was then recorded. **(C)** The cell viability rates were plotted to analyze the data.

We also evaluated the sensitivity of live wild-type *E. coli* cells to VP1∼VP8 and NR. As shown in Fig. 6B-6C and Fig. S5, VP5 showed the best antibacterial activity in a dose-dependent manner, for which the inhibition rates at 200 or 400 μg/mL were ∼50% and ∼97%, respectively. Hence, VP5 shows promise as an antimicrobial peptide against wild type *E. coli*.

### VP5 binds with DnaK through canonical substrate binding pocket in predicted structure

To determine if DnaK binds to VP5 through SBD domain, we performed a blind global protein-peptide docking using HPEPDOCK web server (http://huanglab.phys.hust.edu.cn/hpepdock/). The isolated DnaK-SBD structure (PDB ID: 1DKZ) was used at this step. The docking result which showed the best docking energy score indicated that a coil-like VP5 binds with DnaK-SBD in the canonical substrate binding pocket and has very few extensive interactions with SBDα (Fig. 7A). This result better explained the fast binding kinetics of VP5 with DnaK in the absence of ATP than other predicted structure which shows significant interactions between VP5 and SBDα (Fig. S6). To further verify this docking result, we performed a competitive fluorescence polarization assay (Fig. 7B). Not surprisingly, 400 μM NR or VP5 significantly blocked the binding of NR-FITC with DnaK, furthermore, the well-known mutant DnaK-L_3,4_^14^ (Loop_3,4_ is replaced by MGG) completely disrupted the binding ability of DnaK to VP5, both results further indicated that VP5 should bind to the same pocket of DnaK as NR in the absence of ATP.

**Figure 7.**
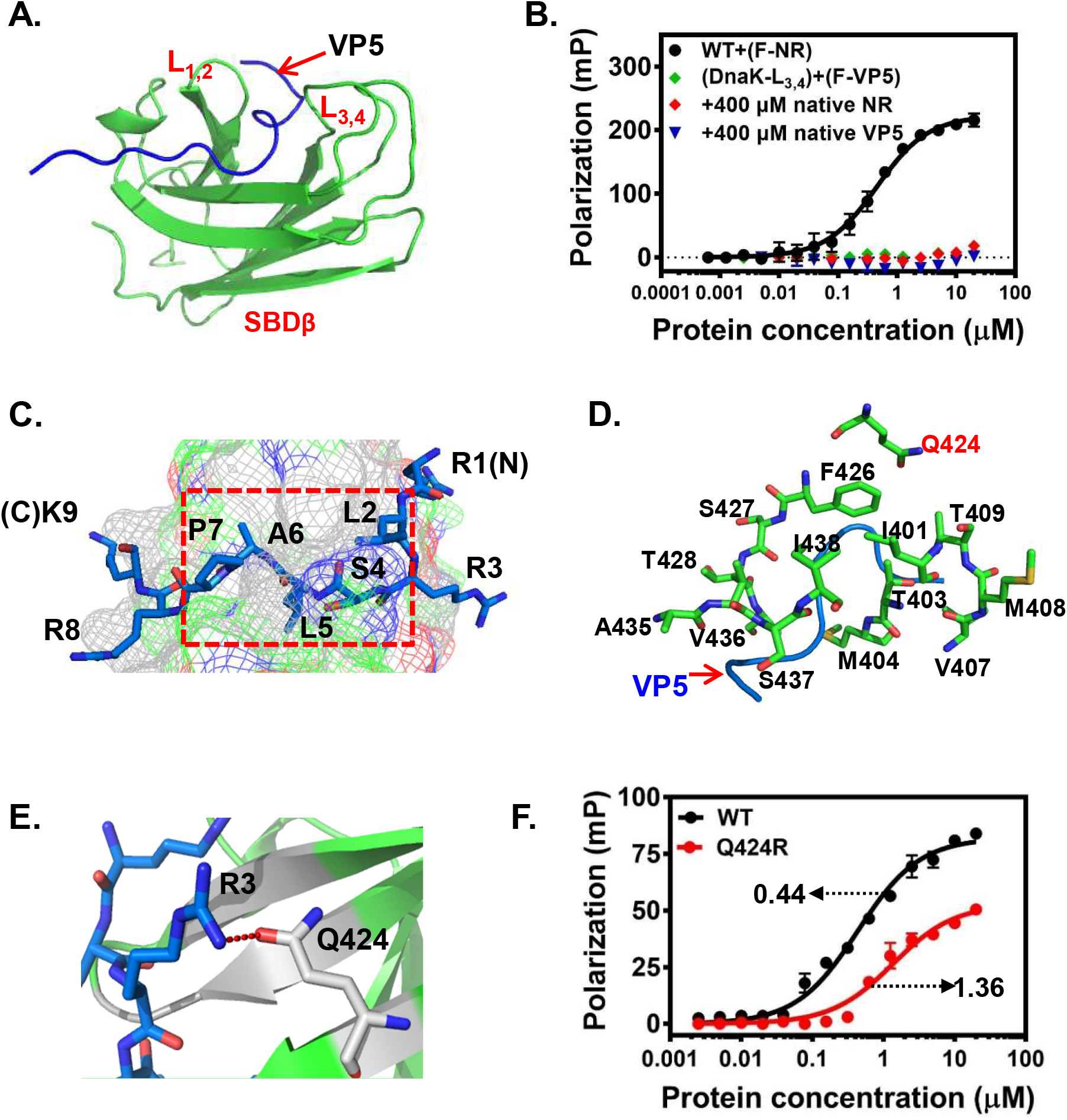
Structure prediction of DnaK binding with VP5. **(A)** A coil-like structure of VP5 can well bind with the substrate binding pocket (composed of L_1,2_ and L_3,4_). Binding interaction between VP5 and DnaK was visualized using PyMOL. VP5 was shown in blue and DnaK SBDβ domain was shown in green. **(B)** Both 400 µM native NR and VP5 could compete the binding of FITC-NR to DnaK; and the Loop_3,4_ mutant (DnaK-L_3,4_) lost the binding activity with FITC-VP5. **(C)** The top view of the substrate binding cleft showing the important interacting residues in VP5 (RLRSLAPRK) and are indicated with blue sticks. **(D)** Closer view to the substrate binding pocket that VP5 binds with. Interacting amino acid residues are shown in green sticks. **(E)** The hydrogen bond between the interaction of Arginine from VP5 and Q424 from β3 of DnaK is shown in red. **(F)** Mutagenesis analysis indicated that VP5 had a reduced binding affinity with mutant DnaK Q424R. The K_*d*_ value (µM) of VP5 binding with each protein was indicated with arrow.

In detail, we found that the N-terminal residues of VP5 mainly participate in the interactions with SBDβ (Fig. 7C), and all hydrophobic residues from N-terminus of VP5 were buried in the substrate binding pocket cleft, leaving the positively charged residue R3 interact with vicinity (Q424 from β3) by additional hydrogen bond, so as to support the high binding affinity of VP5 with SBDβ (Fig. 7D-7E). Our mutagenesis experiment indicated that the binding affinity of DnaK-Q424R to VP5 was obviously lower than that in wide type DnaK (Fig. 7F), further validated our predicted structure as shown in figure 7A.

Unfortunately, no reasonable docking results to reflect how DnaK binds to VP5 in the presence of ATP were acquired by using DnaK-ATP structures (PDB ID: 4JNE & 4B9Q), which needs further examinations in the future.

In conclusion (Fig. 8), we have identified a novel native polypeptide VP5, which might bind DnaK through the canonical substrate binding pocket in a coil-like conformation and not tightly regulated by classical allosteric coupling mechanism. Sequences such as VP5 with enriched positively charged residues are easily exposed to protein surface, DnaK may bind to these polypeptide sequences through divergent mode to exercise other chaperone functions, instead of helping protein (re)folding.

**Figure 8.**
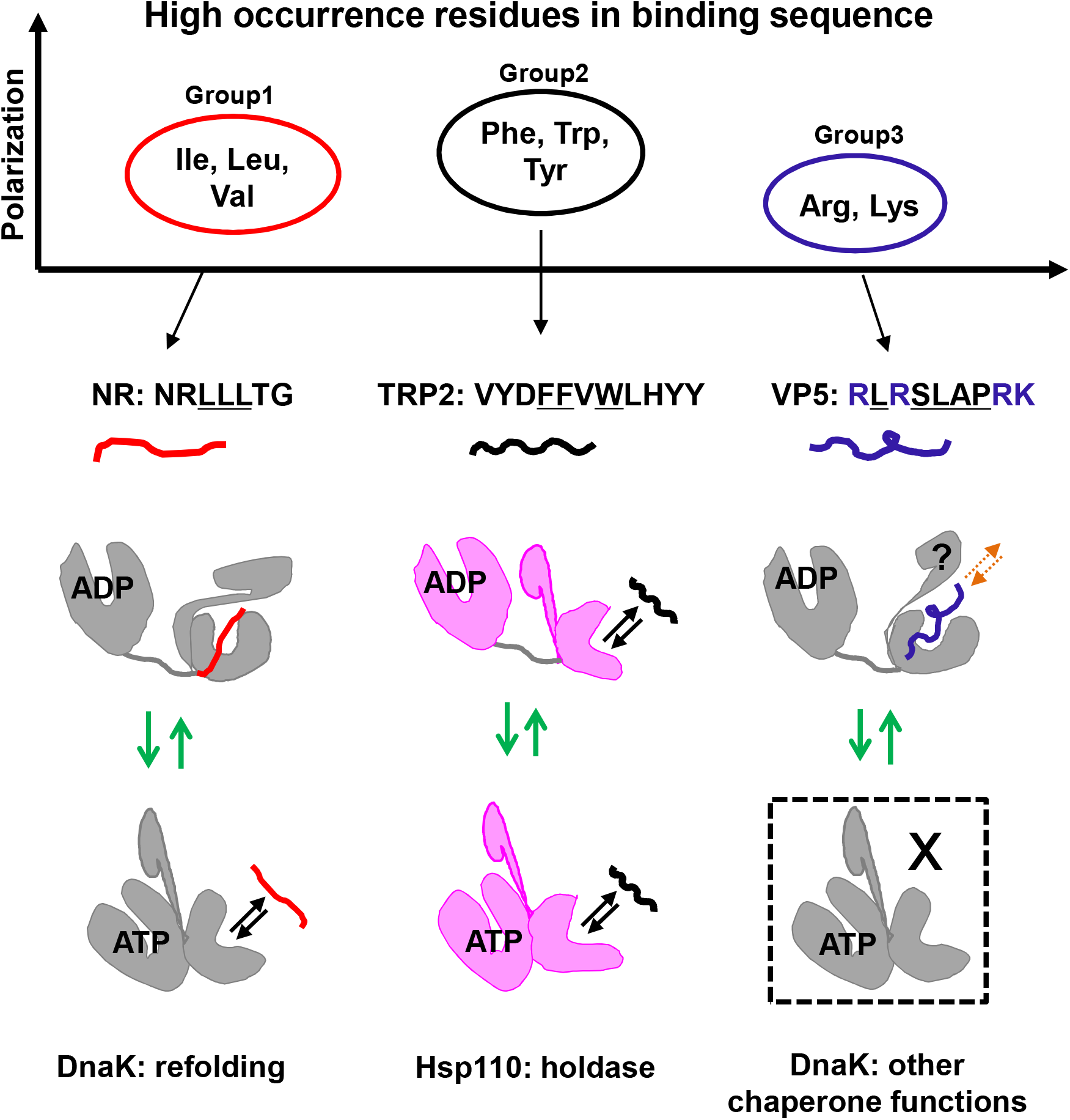
Theoretical binding models for three representative peptide substrates of heat shock proteins. Three different groups of residues are found with very high occurrence in binding sequences of heat shock proteins, they are respectively group 1 aliphatic residues with high percentage of Ile, Leu and Val (as seen in NR: NRLLLTG); group 2 aromatic residues with high percentage of Phe, Trp and Tyr (as seen in TRP2: VYDFFVWLHYY); group 3 basic amino acid residues with high percentage of Arg and Lys (as seen in VP5: RLRSLAPRK). The key residues in each peptide substrate to interact with heat shock proteins were emphasized with underlines; and the basic residues in VP5 were labeled in blue. The peptide substrate types determine their binding mode, thus inducing divergent peptide substrate binding properties.

## Discussion

Hsp70s play a key role in maintaining proteostasis via mediating the proper (re)folding of newly synthesized, misfolded or aggregated proteins.^36^ Although the molecular mechanisms underlying Hsp70’s action have been widely studied and several Hsp70-binding polypeptides have been identified,^24, 27, 33, 37-42^ due to a much more complicated circumstances in living cells, our current knowledge of Hsp70-recognized polypeptide substrates is still limited.

In the present study, we initiated a screening with an iPDMS peptide microarray to systematically study the characteristics of DnaK-binding polypeptides. Unlike previous phage-display or cellulose-bound peptides scan, our peptide microarray screening offers the following advantages: (i) high detection sensitivity owing to the “zero background” nano-membrane of the peptide microarray,^32^ (ii) biased peptide sequence selection from biologically relevant proteins that are either regulated by molecular chaperones (*e*.*g*., native proteins that are involved in autoimmune diseases and infectious diseases) or potentially interact with Hsp70s (*e*.*g*., native proteins that are related with cancer disease, such as Her2), greatly increasing the response rate by ∼17 folds compared with the traditional phase-display method,^33-34^ (iii) high reliability of data sets to exclude randomly responsive polypeptides through multiple-round screenings. Thus, we have set up a superior screening system to obtain DnaK (or other chaperone) interacting peptide substrates in high efficiency by taking the advantage of iPDMS-based microarray.

Not surprisingly, the amino acid distribution among the identified DnaK-binding polypeptides VP1∼VP8 in our study is largely similar to previous findings,^33^ further suggesting the conservatism of DnaK-binding polypeptides. Notably, the occurrence of proline in VP1∼VP8 is very high, though different from the previous finding of cellulose-bound peptides scan,^33^ is consistent with the conclusion that short proline-rich polypeptides have specific tendency to interact with DnaK.^41-44^

Our interesting discovery demonstrated that VP5 presented a non-classical binding characteristics, which validates that the substrate binding properties of Hsp70s in organisms are complex. This conclusion should be easily accepted in the field of molecular chaperone. For instance, Hsp110s, distant homologues of Hsp70s, bind and release peptide substrates only in fast kinetic mode, which is quite different from Hsp70s. ^45-46^ Moreover, recent studies indicated that Hsp70s have two substrate binding sites,^31^ and the substrate binding properties in these two binding sites are also different.

Importantly, we finally claimed that the divergent substrate binding properties are mainly determined by the properties of peptide substrates themselves, and summarized the theoretical working models for hitherto representative types of peptide substrate binding with heat shock proteins by integrating on our data ^14, 19, 45^ (Fig. 8). Our novel polypeptide VP5, with more basic residues, was observed performing unique substrate binding properties in our hand. Based on its medium level substrate binding kinetics with DnaK in the absence of ATP ^14, 19, 31, 45^ (Table S2), we concluded that the conformation of substrate binding pocket lid (SBDα) when VP5 binds with DnaK in the absence of ATP might be in a transition state (Fig. 8), which is between fully open status (as seen in TRP2 binding with Hsp110) and tightly closed status (as seen in NR binding with DnaK), complying with faster binding kinetics but high affinity binding properties. Supporting our hypothesis, we see a coil-like VP5 bind inside of SBDβ (Fig. 7A), making the substrate binding pocket a bit crowded. In such condition, the SBDα should stay away a little bit from SBDβ in order to provide more space for accommodating this bigger client. Of note, this predicted structure might have just provided one possible conformation, and our theoretical working model for VP5 binding with DnaK will be revised as soon as the crystal structure of DnaK/VP5 complex was solved.

Though the inhibitory effect of VP5 on DnaK refolding activity was not as efficient as VP4 *in vitro*, VP5 has superior inhibitory activity on cellular viability over VP4. The peptide sequences which contain more positively charged residues are popularly used as membrane penetrating peptide,^47-48^ thus VP5 can penetrate live cells easier than VP4 to disturb DnaK refolding activity and influence cellular growth. We are confident that VP5 has promising clinical applications as an excellent lead peptide for antibacterial drug development.

## Materials and Methods

### Peptide Microarray

In this study, the polymer coated initiator integrated poly (dimethysiloxane) (iPDMS)-based peptide microarray comprised four different types of polypeptides derived from 78 native proteins. The related information for included native proteins were shown in Table S1.

2645 polypeptides derived from above proteins were synthesized (GENEWIZ, Shanghai, China) to fabricate Microarray 1. Briefly, ∼0.6 nL of each polypeptide with concentration of 0.1 mg/mL was printed onto the activated nanomembranes by contact spotter Smart 48 (Capital Bio, Beijing, China) to form 6×6 array for each well. In each well, there are three positive controls printed with His-tagged DnaK at the concentration of 10 μg/mL and one negative control printed with reaction buffer. Both Microarray 2 and Microarray 3 were fabricated just like Microarray 1, but with downsized capacity of polypeptides.

### The screening of DnaK binding polypeptides with iPDMS-based peptide microarray

To identify DnaK binding polypeptides, DnaK was screened using iPDMS-based peptide microarray as previously described with minor modifications.^49^ DnaK was diluted with buffer A (25 mM HEPES, 150 mM KCl, 10 mM Mg(OAc)_2_, 10% glycerol, 1 mM DTT) to indicated concentrations, then 100 μL DnaK dilution was added to each well and incubated with peptide microarray at 500 rpm for 1 h at room temperature. Subsequently, the microarray was washed extensively with buffer A to remove unbound DnaK and incubated with 100 μL anti-His antibody (HRP conjugated, Abclonal) at 500 rpm for 0.5 h at room temperature. After second washing, redundant anti-His antibody was washed away and 15 μL chemiluminescence substrate (Thermo Scientific) was added onto each well to generate chemiluminescence signal. The images were taken at 635 nm using the Clear 4 imaging system (Suzhou Epitope, China) and analyzed with MATLAB.

### Protein expression and purification

All proteins used in this study were expressed and purified similar to previous descriptions. For His-tagged DnaK, the DnaK construct (residues 2-610) was cloned into pET-21a (+) vector and expressed in BL21 (DE3) with 1 mM IPTG induction at 30 °C for 5 h. After sonication, the His-tagged DnaK protein was purified using a HisTrap HP column (GE Healthcare) with 2×PBS buffer followed by further purification using a HiTrap Q column (GE Healthcare) with 25 mM HEPES-KOH buffer (pH 7.5).

For DnaK (residues 2-610), DnaK-L_3,4_, DnaJ and GrpE, the constructs were cloned into pSMT3 vector and expressed in BL21 (DE3) with 1 mM IPTG induction at 30 °C for 5 h. The Smt3 fusion proteins were first purified on a HisTrap HP column, and dialyzed against the same buffer as used during the first step purification. The protein was then purified on a second HisTrap HP column after digesting with Ulp1 to remove Smt3 tag, and was further purified on a HiTrap Q column or a Superdex 75 10/300 GL column (GE Healthcare). For DnaK mutant proteins, the constructs were generated using QuickMutation(tm) Site-Directed Mutagenesis kit (Beyotime) and the protein expression and purification procedures were extremely the same as wt DnaK protein: DnaK (residues 2-610).

### Fluorescence anisotropy peptide substrate binding assay

The fluorescein-labeled polypeptides were synthesized by GENEWIZ. To examine the binding affinities for different polypeptides, serial dilutions of DnaK protein were premixed with fluorescein-labeled polypeptides at ideal concentration in 384-well plate at room temperature, either in the absence of or presence of ATP. After reaching equilibrium, the fluorescence anisotropy was measured on Cytation 3 multi-mode reader (BioTek). The data were fitted to one-site specific binding equation using Prism GraphPad to calculate dissociation constants (K*d*). Experiments were conducted at least three times with two batches of proteins.

### Peptide substrate binding kinetics measurements

The binding kinetics of each polypeptide were measured by mixing fluorescein-labeled polypeptides with 10 µM DnaK protein. Polypeptides alone were used for initial readings. After adding DnaK protein, readings were recorded every 10 or 30 s. For the measurement in the presence of ATP, DnaK protein was first incubated with 2 mM ATP for 2 min before adding fluorescein-labeled polypeptides.

### Luciferase refolding assay

Firefly luciferase (Promega) was diluted with buffer B (25 mM HEPES-KOH, pH 7.5, 10 mM Mg (OAc)_2_, 100 mM KCl, 3 mM ATP and 1 mM DTT) to a final concentration of 0.2 μM in the presence of 3 μM DnaK. After heating at 42 °C for 15 min, refolding reactions were carried out by adding the heated luciferase into a reaction system containing 3 μM DnaK, 0.67 μM DnaJ, and 0.33 μM GrpE with or without tested polypeptides. After incubating for 30 min, luciferase activities were measured in Cytation 3 multi-mode reader (BioTek) by quickly mixing 2 μL of above refolding mixture with 50 μL of luciferase substrate (Promega). The activity of unheated luciferase in buffer B was set as 100%.

### Determination of antibacterial activities

Wild-type (wt) *E. coli* (ATCC 25922) were grown overnight in LB broth and seeded into 96-well plate. Serially diluted stock solutions of polypeptides were then added to each well prior to overnight growth at 42 °C for ∼18 h. The OD_600_ of each well was then recorded with Cytation 3 multi-mode reader (BioTek). The well of cells without any polypeptides was set as the positive control.

## Acknowledgments

We are grateful to Dr. Qinglian Liu and Dr. Yiming Ma for providing insightful suggestions to our study. This study was supported by Jiangsu Natural Science Foundation (BK20190230) and National Natural Science Foundation of China (31901065).

## Author Contributions

J.Y. designed the study; Y.S. and H.Z. performed all the experiments and analyzed the data; J.Y. wrote the manuscript. All the authors contributed to the revision and review of the manuscript.

**Supplementary table 1**. Native protein sources for iPDMS-based peptide microarray.

**Supplementary table 2**. The comparisons of four different peptide substrate binding properties.

**Supplementary Figure 1**. Polypeptides VP1∼VP8 showed binding activity to DnaK.

**Supplementary figure 2**. Substrate binding kinetics of polypeptides with DnaK in the absence of and presence of ATP.

**Supplementary figure 3**. ATP mediates the release of different polypeptides from DnaK.

**Supplementary figure 4**. The identified polypeptides inhibit the refolding activity of DnaK.

**Supplementary figure 5**. VP5 inhibits the viability of wild-type *E. coli* cells more efficiently than other identified polypeptides.

**Supplementary figure 6**. The predicted VP5 interacting residues from SBD.

## References

1. Dissmeyer, N.; Coux, O.; Rodriguez, M. S.; Barrio, R.; Core Group Members of, P., PROTEOSTASIS: A European Network to Break Barriers and Integrate Science on Protein Homeostasis. Trends Biochem Sci 2019, 44 (5), 383–387.

2. Kim, Y. E.; Hipp, M. S.; Bracher, A.; Hayer-Hartl, M.; Hartl, F. U., Molecular chaperone functions in protein folding and proteostasis. Annu Rev Biochem 2013, 82, 323–55.

3. Fernandez-Fernandez, M. R.; Valpuesta, J. M., Hsp70 chaperone: a master player in protein homeostasis. F1000Res 2018, 7.

4. Cappucci, U.; Noro, F.; Casale, A. M.; Fanti, L.; Berloco, M.; Alagia, A. A.; Grassi, L.; Le Pera, L.; Piacentini, L.; Pimpinelli, S., The Hsp70 chaperone is a major player in stress-induced transposable element activation. Proc Natl Acad Sci U S A 2019, 116 (36), 17943–17950.

5. Rosenzweig, R.; Nillegoda, N. B.; Mayer, M. P.; Bukau, B., The Hsp70 chaperone network. Nat Rev Mol Cell Biol 2019, 20 (11), 665–680.

6. Evans, C. G.; Chang, L.; Gestwicki, J. E., Heat shock protein 70 (hsp70) as an emerging drug target. J Med Chem 2010, 53 (12), 4585–602.

7. Garbuz, D. G.; Zatsepina, O. G.; Evgen’ev, M. B., [The Major Human Stress Protein Hsp70 as a Factor of Protein Homeostasis and a Cytokine-Like Regulator]. Mol Biol (Mosk) 2019, 53 (2), 200–217.

8. Arlet, J. B.; Ribeil, J. A.; Guillem, F.; Negre, O.; Hazoume, A.; Marcion, G.; Beuzard, Y.; Dussiot, M.; Moura, I. C.; Demarest, S.; de Beauchene, I. C.; Belaid-Choucair, Z.; Sevin, M.; Maciel, T. T.; Auclair, C.; Leboulch, P.; Chretien, S.; Tchertanov, L.; Baudin-Creuza, V.; Seigneuric, R.; Fontenay, M.; Garrido, C.; Hermine, O.; Courtois, G., HSP70 sequestration by free alpha-globin promotes ineffective erythropoiesis in beta-thalassaemia. Nature 2014, 514 (7521), 242–6.

9. Yang, S.; Xiao, H.; Cao, L., Recent advances in heat shock proteins in cancer diagnosis, prognosis, metabolism and treatment. Biomed Pharmacother 2021, 142, 112074.

10. Rutledge, B. S.; Choy, W. Y.; Duennwald, M. L., Folding or holding?-Hsp70 and Hsp90 chaperoning of misfolded proteins in neurodegenerative disease. J Biol Chem 2022, 298 (5), 101905.

11. Lubkowska, A.; Pluta, W.; Stronska, A.; Lalko, A., Role of Heat Shock Proteins (HSP70 and HSP90) in Viral Infection. Int J Mol Sci 2021, 22 (17).

12. Ferraro, M.; D’Annessa, I.; Moroni, E.; Morra, G.; Paladino, A.; Rinaldi, S.; Compostella, F.; Colombo, G., Allosteric Modulators of HSP90 and HSP70: Dynamics Meets Function through Structure-Based Drug Design. J Med Chem 2019, 62 (1), 60–87.

13. Yang, J.; Nune, M.; Zong, Y.; Zhou, L.; Liu, Q., Close and Allosteric Opening of the Polypeptide-Binding Site in a Human Hsp70 Chaperone BiP. Structure 2015, 23 (12), 2191–2203.

14. Qi, R.; Sarbeng, E. B.; Liu, Q.; Le, K. Q.; Xu, X.; Xu, H.; Yang, J.; Wong, J. L.; Vorvis, C.; Hendrickson, W. A.; Zhou, L.; Liu, Q., Allosteric opening of the polypeptide-binding site when an Hsp70 binds ATP. Nat Struct Mol Biol 2013, 20 (7), 900–7.

15. Yang, J.; Zong, Y.; Su, J.; Li, H.; Zhu, H.; Columbus, L.; Zhou, L.; Liu, Q., Conformation transitions of the polypeptide-binding pocket support an active substrate release from Hsp70s. Nat Commun 2017, 8 (1), 1201.

16. Zhuravleva, A.; Gierasch, L. M., Substrate-binding domain conformational dynamics mediate Hsp70 allostery. Proc Natl Acad Sci U S A 2015, 112 (22), E2865–73.

17. Mayer, M. P., Hsp70 chaperone dynamics and molecular mechanism. Trends Biochem Sci 2013, 38 (10), 507–14.

18. Kityk, R.; Kopp, J.; Sinning, I.; Mayer, M. P., Structure and dynamics of the ATP-bound open conformation of Hsp70 chaperones. Mol Cell 2012, 48 (6), 863–74.

19. Zhu, X.; Zhao, X.; Burkholder, W. F.; Gragerov, A.; Ogata, C. M.; Gottesman, M. E.; Hendrickson, W. A., Structural analysis of substrate binding by the molecular chaperone DnaK. Science 1996, 272 (5268), 1606–14.

20. Jordan, R.; McMacken, R., Modulation of the ATPase activity of the molecular chaperone DnaK by peptides and the DnaJ and GrpE heat shock proteins. J Biol Chem 1995, 270 (9), 4563–9.

21. Piette, B. L.; Alerasool, N.; Lin, Z. Y.; Lacoste, J.; Lam, M. H. Y.; Qian, W. W.; Tran, S.; Larsen, B.; Campos, E.; Peng, J.; Gingras, A. C.; Taipale, M., Comprehensive interactome profiling of the human Hsp70 network highlights functional differentiation of J domains. Mol Cell 2021, 81 (12), 2549–2565 e8.

22. Bracher, A.; Verghese, J., The nucleotide exchange factors of Hsp70 molecular chaperones. Front Mol Biosci 2015, 2, 10.

23. Marcinowski, M.; Holler, M.; Feige, M. J.; Baerend, D.; Lamb, D. C.; Buchner, J., Substrate discrimination of the chaperone BiP by autonomous and cochaperone-regulated conformational transitions. Nat Struct Mol Biol 2011, 18 (2), 150–8.

24. Romanova, E. A.; Sharapova, T. N.; Telegin, G. B.; Minakov, A. N.; Chernov, A. S.; Ivanova, O. K.; Bychkov, M. L.; Sashchenko, L. P.; Yashin, D. V., A 12-mer Peptide of Tag7 (PGLYRP1) Forms a Cytotoxic Complex with Hsp70 and Inhibits TNF-Alpha Induced Cell Death. Cells 2020, 9 (2).

25. Albakova, Z.; Armeev, G. A.; Kanevskiy, L. M.; Kovalenko, E. I.; Sapozhnikov, A. M., HSP70 Multi-Functionality in Cancer. Cells 2020, 9 (3).

26. Dalphin, M. D.; Stangl, A. J.; Liu, Y.; Cavagnero, S., KLR-70: A Novel Cationic Inhibitor of the Bacterial Hsp70 Chaperone. Biochemistry 2020, 59 (20), 1946–1960.

27. Stevens, S. Y.; Cai, S.; Pellecchia, M.; Zuiderweg, E. R., The solution structure of the bacterial HSP70 chaperone protein domain DnaK(393-507) in complex with the peptide NRLLLTG. Protein Sci 2003, 12 (11), 2588–96.

28. Wang, W.; Liu, Q.; Liu, Q.; Hendrickson, W. A., Conformational equilibria in allosteric control of Hsp70 chaperones. Mol Cell 2021, 81 (19), 3919–3933 e7.

29. Lu, J.; Zhang, X.; Wu, Y.; Sheng, Y.; Li, W.; Wang, W., Energy landscape remodeling mechanism of Hsp70-chaperone-accelerated protein folding. Biophys J 2021, 120 (10), 1971–1983.

30. Wang, W.; Hendrickson, W. A., Intermediates in allosteric equilibria of DnaK-ATP interactions with substrate peptides. Acta Crystallogr D Struct Biol 2021, 77 (Pt 5), 606–617.

31. Li, H.; Zhu, H.; Sarbeng, E. B.; Liu, Q.; Tian, X.; Yang, Y.; Lyons, C.; Zhou, L.; Liu, Q., An unexpected second binding site for polypeptide substrates is essential for Hsp70 chaperone activity. J Biol Chem 2020, 295 (2), 584–596.

32. Ma, H.; Wu, Y.; Yang, X.; Liu, X.; He, J.; Fu, L.; Wang, J.; Xu, H.; Shi, Y.; Zhong, R., Integrated poly(dimethysiloxane) with an intrinsic nonfouling property approaching “absolute” zero background in immunoassays. Anal Chem 2010, 82 (15), 6338–42.

33. Rudiger, S.; Germeroth, L.; Schneider-Mergener, J.; Bukau, B., Substrate specificity of the DnaK chaperone determined by screening cellulose-bound peptide libraries. EMBO J 1997, 16 (7), 1501–7.

34. Gragerov, A.; Zeng, L.; Zhao, X.; Burkholder, W.; Gottesman, M. E., Specificity of DnaK-peptide binding. J Mol Biol 1994, 235 (3), 848–54.

35. Nordquist, E. B.; English, C. A.; Clerico, E. M.; Sherman, W.; Gierasch, L. M.; Chen, J., Physics-based modeling provides predictive understanding of selectively promiscuous substrate binding by Hsp70 chaperones. PLoS Comput Biol 2021, 17 (11), e1009567.

36. Mayer, M. P.; Bukau, B., Hsp70 chaperones: cellular functions and molecular mechanism. Cell Mol Life Sci 2005, 62 (6), 670–84.

37. Lam, K. T.; Calderwood, S. K., hsp70 binds specifically to a peptide derived from the highly conserved domain (I) region of p53. Biochem Biophys Res Commun 1992, 184 (1), 167–74.

38. Ivey, R. A., 3rd; Subramanian, C.; Bruce, B. D., Identification of a Hsp70 recognition domain within the rubisco small subunit transit peptide. Plant Physiol 2000, 122 (4), 1289–99.

39. Okochi, M.; Hayashi, H.; Ito, A.; Kato, R.; Tamura, Y.; Sato, N.; Honda, H., Identification of HLA-A24-restricted epitopes with high affinities to Hsp70 using peptide arrays. J Biosci Bioeng 2008, 105 (3), 198–203.

40. Stocki, P.; Wang, X. N.; Morris, N. J.; Dickinson, A. M., HSP70 natively and specifically associates with an N-terminal dermcidin-derived peptide that contains an HLA-A*03 antigenic epitope. J Biol Chem 2011, 286 (14), 12803–11.

41. Czihal, P.; Knappe, D.; Fritsche, S.; Zahn, M.; Berthold, N.; Piantavigna, S.; Muller, U.; Van Dorpe, S.; Herth, N.; Binas, A.; Kohler, G.; De Spiegeleer, B.; Martin, L. L.; Nolte, O.; Strater, N.; Alber, G.; Hoffmann, R., Api88 is a novel antibacterial designer peptide to treat systemic infections with multidrug-resistant Gram-negative pathogens. ACS Chem Biol 2012, 7 (7), 1281–91.

42. Zahn, M.; Kieslich, B.; Berthold, N.; Knappe, D.; Hoffmann, R.; Strater, N., Structural identification of DnaK binding sites within bovine and sheep bactenecin Bac7. Protein Pept Lett 2014, 21 (4), 407–12.

43. Knappe, D.; Zahn, M.; Sauer, U.; Schiffer, G.; Strater, N.; Hoffmann, R., Rational design of oncocin derivatives with superior protease stabilities and antibacterial activities based on the high-resolution structure of the oncocin-DnaK complex. Chembiochem 2011, 12 (6), 874–6.

44. Zahn, M.; Berthold, N.; Kieslich, B.; Knappe, D.; Hoffmann, R.; Strater, N., Structural studies on the forward and reverse binding modes of peptides to the chaperone DnaK. J Mol Biol 2013, 425 (14), 2463–79.

45. Xu, X.; Sarbeng, E. B.; Vorvis, C.; Kumar, D. P.; Zhou, L.; Liu, Q., Unique peptide substrate binding properties of 110-kDa heat-shock protein (Hsp110) determine its distinct chaperone activity. J Biol Chem 2012, 287 (8), 5661–72.

46. Li, H.; Hu, L.; Cuffee, C. W.; Mohamed, M.; Li, Q.; Liu, Q.; Zhou, L.; Liu, Q., Interdomain interactions dictate the function of the Candida albicans Hsp110 protein Msi3. J Biol Chem 2021, 297 (3), 101082.

47. Castelletto, V.; Barnes, R. H.; Karatzas, K. A.; Edwards-Gayle, C. J. C.; Greco, F.; Hamley, I. W.; Rambo, R.; Seitsonen, J.; Ruokolainen, J., Arginine-Containing Surfactant-Like Peptides: Interaction with Lipid Membranes and Antimicrobial Activity. Biomacromolecules 2018, 19 (7), 2782–2794.

48. Kanwar, J. R.; Gibbons, J.; Verma, A. K.; Kanwar, R. K., Cell-penetrating properties of the transactivator of transcription and polyarginine (R9) peptides, their conjugative effect on nanoparticles and the prospect of conjugation with arsenic trioxide. Anticancer Drugs 2012, 23 (5), 471–82.

49. Lu, Y.; Li, Z.; Teng, H.; Xu, H.; Qi, S.; He, J.; Gu, D.; Chen, Q.; Ma, H., Chimeric peptide constructs comprising linear B-cell epitopes: application to the serodiagnosis of infectious diseases. Sci Rep 2015, 5, 13364.

