## supplementary tables and supplementary figures for "Conserved and divergent peptide substrate binding properties of bacterial Hsp70"

**Supplementary table 1. Native protein sources for iPDMS-based peptide microarray**

| Microarray Category | Protein source | SUM. (Antigen) | Peptide Counts |
|---|---|---|---|
| Viral Protein | Newcastle Disease Virus (NDV) | 4 | 196 |
| | Avian Influenza Virus (AIV) | 5 | 135 |
| | Peste des petits ruminants Virus (PPRV) | 4 | 197 |
| | Pseudorabies virus (PRV) | 6 | 324 |
| | EV71 | 11 | 208 |
| | SARS-CoV-2 | 4 | 99 |
| Infectious disease related | Mycobacterium tuberculosis (TB) | 12 | 298 |
| | Malaria | 10 | 398 |
| Cancer related | Tumor necrosis factor α (TNF-α) | 1 | 16 |
| | Claudin 18.2 (CLDN18.2) | 1 | 32 |
| | Human epidermal growth factor receptor (Her2) | 1 | 9 |
| | Interleukin-4 receptor (IL-4R) | 1 | 11 |
| | CD20 | 1 | 15 |
| Endogenous Protein (Autoimmune disease) | Type 1 Diabetes Mellitus (T1DM) | 9 | 272 |
| | Autoimmune Thyroid Disease (AITD) | 3 | 229 |
| | Systemic Vasculitis (SV) | 5 | 206 |
| **Total** | | | **2645** |

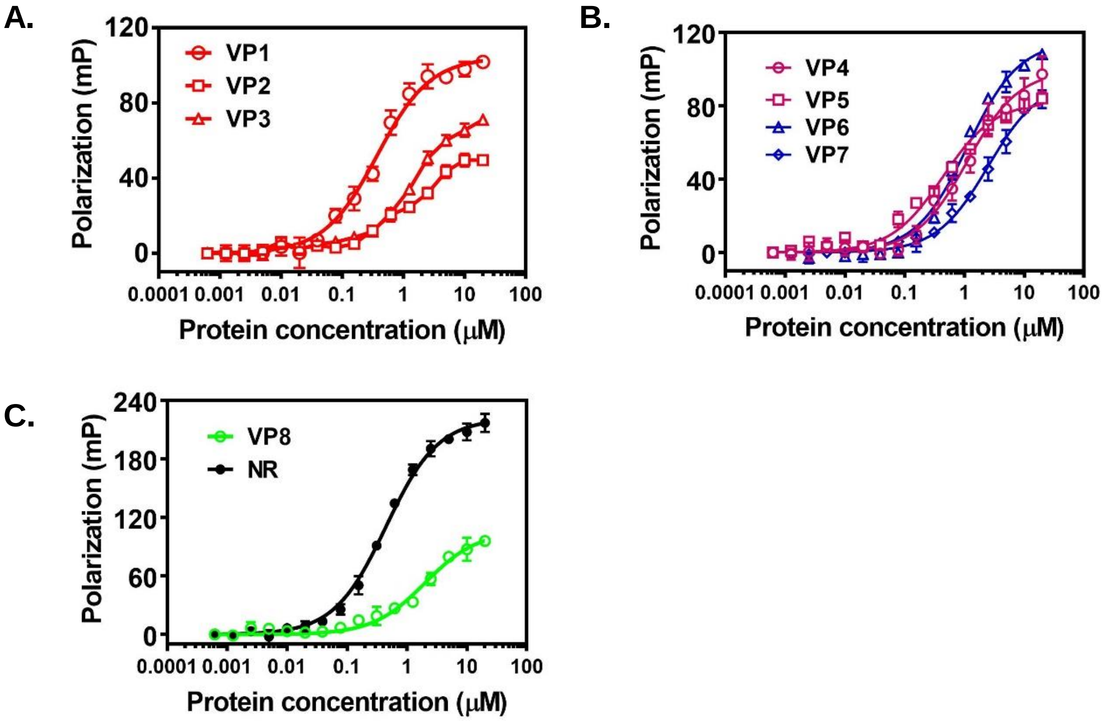


**Supplementary figure 1. Polypeptides VP1~VP8 showed binding activity to DnaK.** To evaluate the binding affinity of VP1~VP8 to DnaK, a fluorescence polarization assay was performed to detect peptide substrate binding. **(A)** VP1, VP2, VP3 ; **(B)** VP4, VP5, VP6, VP7 ; **(C)** VP8 and NR, all bind to DnaK in a dose-dependent manner.

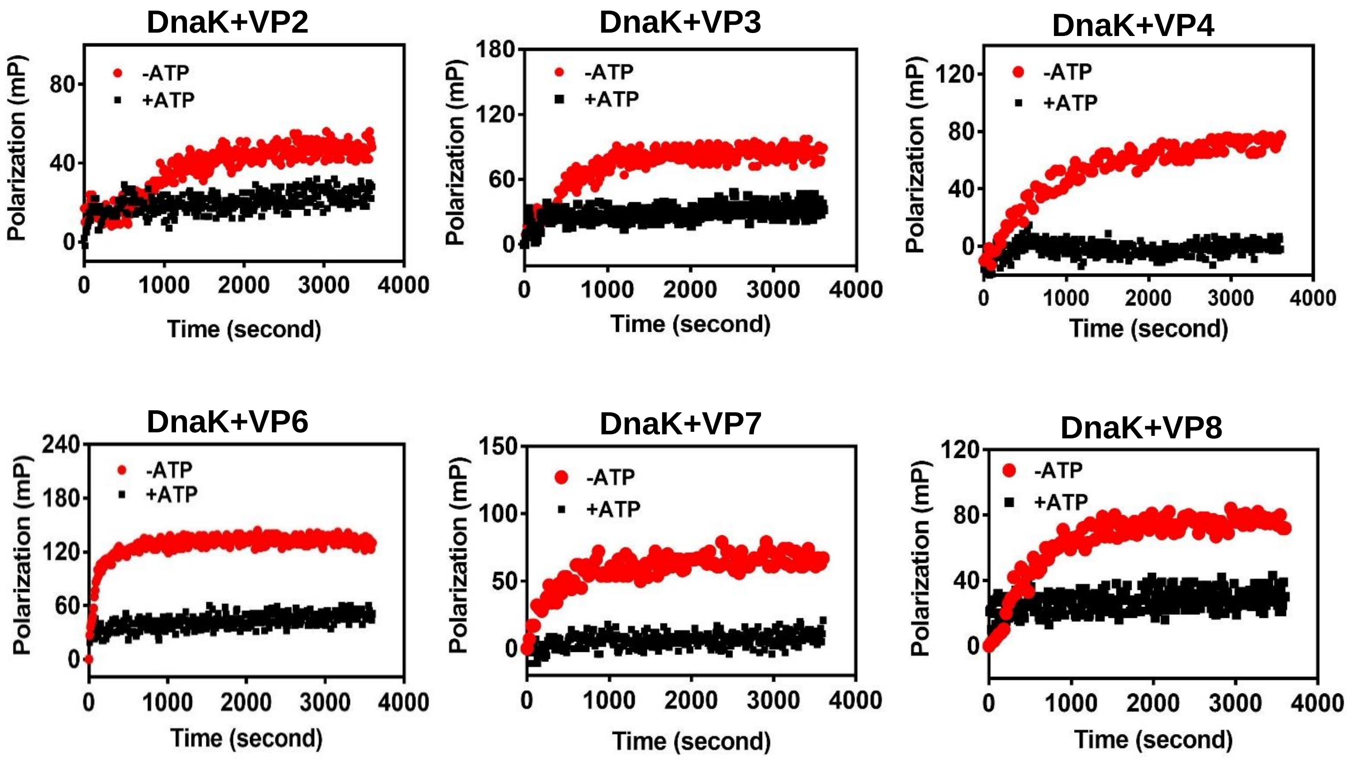


**Supplementary figure 2. Substrate binding kinetics of polypeptides with DnaK in the absence of and presence of ATP.** The fluorescence FITC labeled polypeptide VP2/3/4/6/7/8 was respectively mixed with DnaK (10 μM) to initiate binding, and binding kinetics was recorded immediately as a function of time. To test the effect of ATP on kinetics, DnaK was previously incubated with ATP for 2 minutes before the addition of FITC-labeled polypeptides.

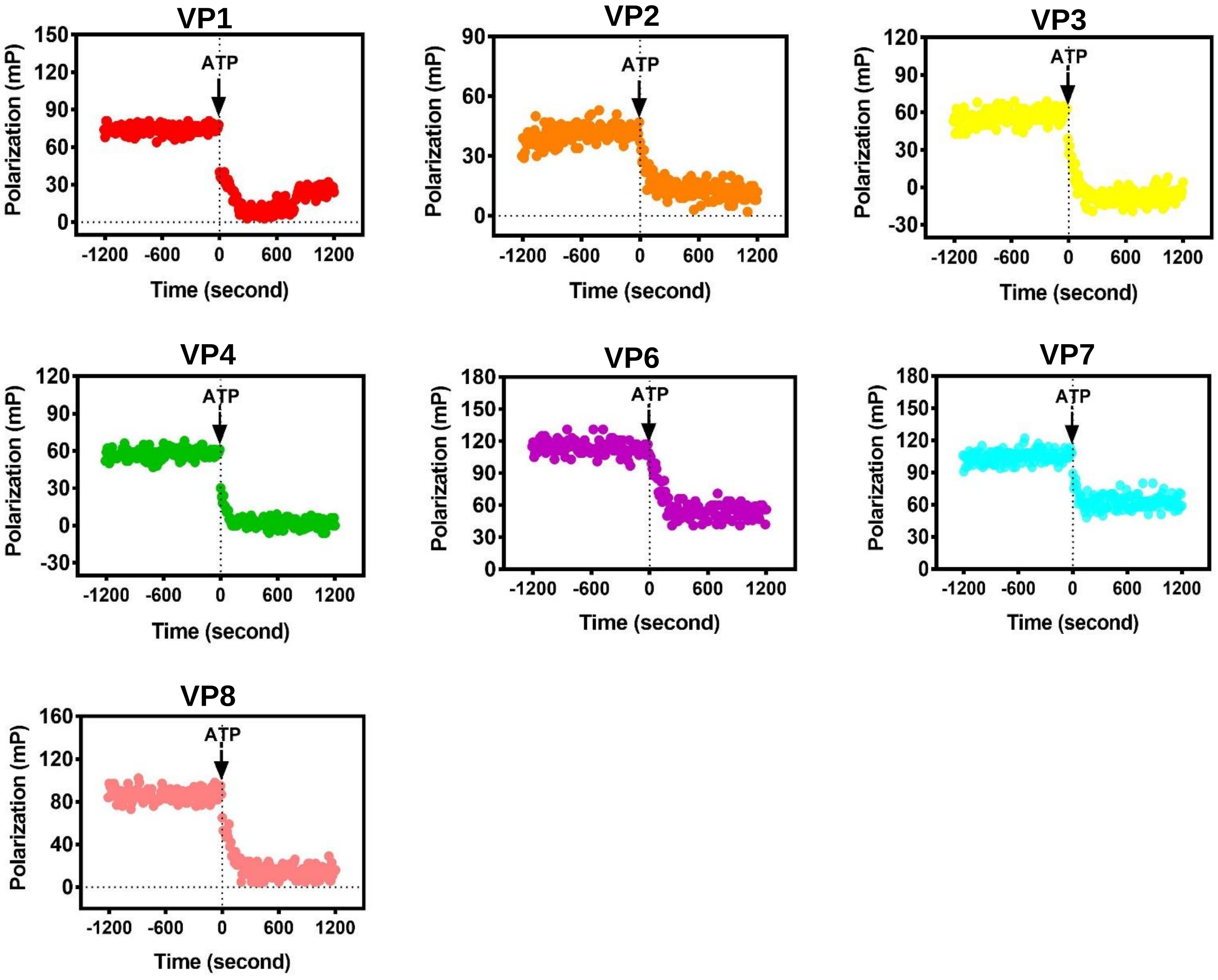


**Supplementary figure 3. ATP mediates the release of different polypeptides from DnaK.** To evaluate if ATP-induced allosteric coupling will induce the release of identified polypeptides, 10 μM DnaK was previously incubated with FITC-labeled VP1/2/3/4/6/7/8 for appropriate time in order to reach for binding equilibrium, then the polarization was traced for 1200 s. By then, 2 mM ATP was added to induce the peptide release, and the dynamic changes of polarization were monitored for another 1200 s. ATP could mediate the release of polypeptides (VP1/2/3/4/6/7/8) from DnaK.

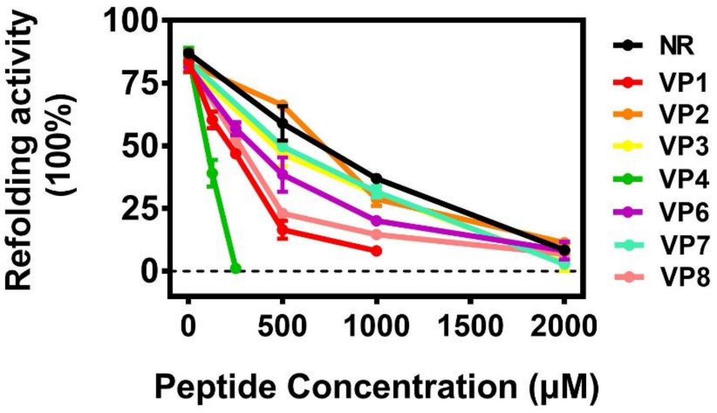


**Supplementary figure 4. The identified polypeptides inhibit the refolding activity of DnaK.** The refolding activities of DnaK were measured in the presence of the indicated concentrations of polypeptides (0, 125, 250, 500, 1000, 2000 μM). VP4 (green line) showed the most efficient inhibitory effect on DnaK refolding activity.

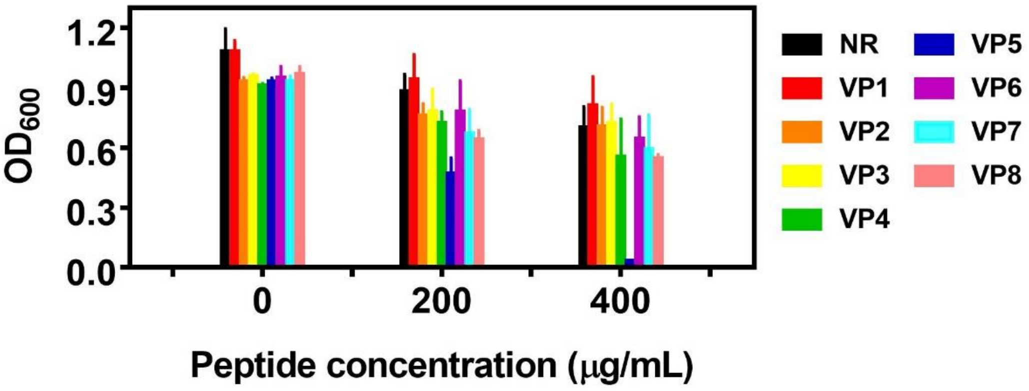


**Supplementary figure 5. VP5 inhibits the viability of wild-type *E. coli* cells more efficiently than other identified polypeptides.** Wild-type *E. coli* (ATCC 25922) cells were seeded into 96-well plate. Two diluted stock solutions of the polypeptides were added in each well prior to overnight growth for ∼18 h at 42℃ respectively, the $OD_{600}$ of each well was then recorded.

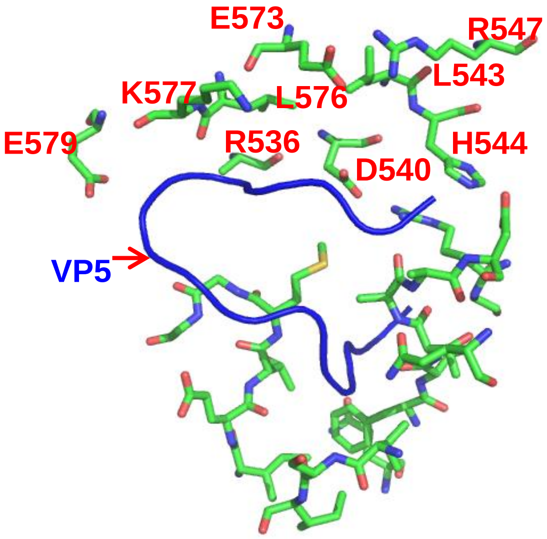


**Supplementary figure 6. The predicted VP5 interacting residues from SBD.** The structure of DnaK-SBD with VP5 complex was predicted by using HPEPDOCK web server. In this structure, the C-terminal residues of VP5 mainly participate in the interactions with SBDβ, leaving N-terminal residues form intensive interactions with SBDα. The VP5 interacting residues from SBD were presented in green sticks, and the interacting residues from SBDα were highlighted in red.

**Supplementary table 2. The comparisons of four different peptide substrate binding properties**

| | (-)ATP | (+)ATP |
|---|---|---|
| **Hsp110-TRP2** | High A<br>Fast K | Low A<br>Fast K |
| **DnaK-NR** | High A<br>Slow K | Low A<br>Fast K |
| **DnaK-TRP2** | Low A<br>Fast K | Low A<br>Fast K |
| **DnaK-VP5** | High A<br>Medium level K | High A<br>Medium level K |

A: binding affinity; K: binding kinetics
